# Analysing protein post-translational modform regions by linear programming

**DOI:** 10.1101/456640

**Authors:** Deepesh Agarwal, Ryan T. Fellers, Bryan P. Early, Dan Lu, Caroline J. DeHart, Philip D. Compton, Paul M. Thomas, Galit Lahav, Neil L. Kelleher, Jeremy Gunawardena

## Abstract

Post-translational modifications (PTMs) at multiple sites can collectively influence protein function but the scope of such PTM coding has been challenging to determine. The number of potential combinatorial patterns of PTMs on a single molecule increases exponentially with the number of modification sites and a population of molecules exhibits a distribution of such “modforms”. Estimating these “modform distributions” is central to understanding how PTMs influence protein function. Although mass-spectrometry (MS) has made modforms more accessible, we have previously shown that current MS technology cannot recover the modform distribution of heavily modified proteins. However, MS data yield linear equations for modform amounts, which constrain the distribution within a high-dimensional, polyhedral “modform region”. Here, we show that linear programming (LP) can efficiently determine a range within which each modform value must lie, thereby approximating the modform region. We use this method on simulated data for mitogen-activated protein kinase 1 with the 7 phosphorylations reported on UniProt, giving a modform region in a 128 dimensional space. The exact dimension of the region is determined by the number of linearly independent equations but its size and shape depend on the data. The average modform range, which is a measure of size, reduces when data from bottom-up (BU) MS, in which proteins are first digested into peptides, is combined with data from top-down (TD) MS, in which whole proteins are analysed. Furthermore, when the modform distribution is structured, as might be expected of real distributions, the modform region for BU and TD combined has a more intricate polyhedral shape and is substantially more constrained than that of a random distribution. These results give the first insights into high-dimensional modform regions and confirm that fast LP methods can be used to analyse them. We discuss the problems of using modform regions with real data, when the actual modform distribution will not be known.

## INTRODUCTION

Post-translational modifications are ubiquitous on most proteins and greatly increase the number of “proteoforms” which participate in cellular processes [1]. Certain modifications require carrier molecules which donate the modifying moiety and enzymes to regulate both forward modification and reverse demodification [19, 27]. Phosphorylation on S, T or Y residues, for example, requires ATP as the carrier molecule and enzymatic regulation is undertaken by protein kinases and phospho-protein phosphatases. Background cellular processes maintain the concentrations of carrier molecules, thereby acting like a chemical battery to drive the modifying reactions. This energy-dissipating architecture confers a distinctive regulatory capability on such reversible, enzymatically-regulated post-translational modifications (hereafter, “PTMs”) [19], on which we will focus in this paper.

Proteins are often modified on multiple amino-acid residues (sites) as well as by different kinds of PTMs. For instance, the transcription factor and “guardian of the genome”, p53, is known to be modified on over 100 sites [4]. If these PTMs were all binary modifications, which would either be present or absent, then the number of potential combinatorial patterns of modification on a single p53 molecule is 2^100^ ≈ 1.3 × 10^30^. This illustrates the extraordinary PTM complexity that can surround even a single cellular protein. Of course, not all these patterns of modification can be present at any one time but that only begs the question of which patterns are present and what their functions are.

Since PTMs effectively replace amino acids by different chemical residues, it is not surprising that they influence protein function. It is not just PTM at a single site but also the pattern of PTMs across an entire protein molecule which can modulate what that molecule does. There is now evidence from many biological contexts of extensive crosstalk between different modified sites [9, 14, 20, 25, 7]. This has suggested the existence of PTM “codes” [2, 13, 26, 15, 17, 6, 16, 8, 24, 11]. The histone code is the best known [8, 24] but p53 itself exhibits complex PTM “barcodes” which determine its varied responses in different cellular circumstances [15]. In this conceptual picture, upstream enzymes “write” and “erase” modifications on a target protein to create “codewords”, which are subsequently “read” by downstream processes. While this idea of information encoding is attractive [19], it has been challenging to confirm the biochemical details in any context. In view of the key role played by PTMs in so many cellular processes, clarifying how PTMs process information has become a central problem of systems biology.

We have previously introduced a quantitative language for analysing this problem [18, 19]. We refer to a combinatorial pattern of PTMs across a single protein molecule as a “modform”. As noted above, the number of potential modforms increases exponentially with the number of modification sites. A given protein will be present within a cell as a population of single molecules and each molecule can, in principle, exhibit its own modform. The most comprehensive measure of the protein’s PTM state is therefore given by the abundance of each modform in the population, which we call the “modform distribution”. This can be thought of as a histogram over the modforms or as a point in a high-dimensional space, in which each dimension, or coordinate axis, corresponds to a specific modform (Fig. 1). If we are to determine how information is encoded by PTMs, then estimating a protein’s modform distribution, at a given time and in a given biological context, is essential. This is the main concern of the present paper.

**Figure 1:**
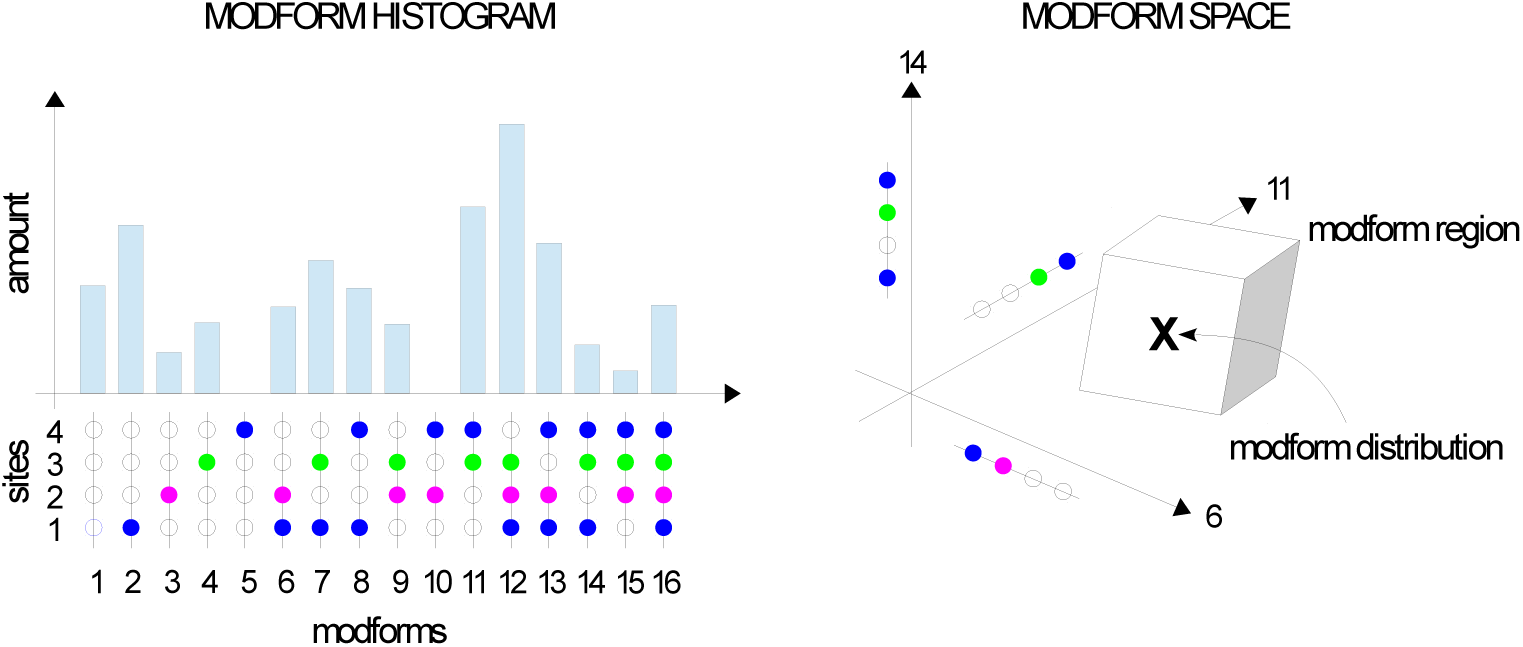
Modforms, distributions and regions. A hypothetical modform distribution is shown as a histogram (left). The protein has 3 types of PTM (blue, magenta, green) at 4 sites, giving 16 modforms in total. The modform distribution can also be viewed as a point (X) in a sixteen-dimensional space (right), where only the three dimensions corresponding to modforms 6, 11 and 14 are shown. Mass-spectrometry data give rise to linear equations which constrain the modform distribution to lie within a modform region (box).

There are a limited number of methods for measuring PTMs. Modification-specific antibodies have been of great importance and have unrivalled sensitivity, including at single-cell level through immunostaining. However, at best, they can only detect PTMs on nearly adjacent sites and are oblivious to the overall modform. Moreover, in comparison to other methods, their quantitative accuracy is suspect [18]. Nuclear magnetic resonance spectroscopy (NMR) is highly quantitative and can reveal certain modform features as well as interactions with binding partners [10, 18] but the limitation to bulk in-vitro measurements has only recently been lifted [12]. Mass spectrometry (MS) remains, at present, the method of choice for estimating modform distributions [18].

In the most-widely used “bottom-up” MS (BU MS), proteins are first proteolytically cleaved into peptides before chromatographic separation and mass determination [21]. So-called “middle-down” MS (MD MS) uses fewer cleavages and correspondingly larger peptides [23]. Peptide modforms can be partly resolved during chromatography and further determined by rounds of fragmentation (MS*^n^*) in the spectrometer, allowing peptide modform distributions to be estimated. However, cleavage severs correlations between modforms on different peptides, leaving the protein modform distribution undetermined [18]. It has seemed conceivable that with multiple proteases with different cleavage patterns, it might still be feasible to reconstruct the protein modform distribution. However, we recently showed mathematically that this is impossible, no matter how many cleavage patterns and proteases are available and that, furthermore, the shortfall in information required to determine the modform distribution increases exponentially with the number of modification sites [3].

Although not yet so widely used, MS can now be undertaken on an intact protein by “top-down” MS (TD MS), which maintains correlations across the protein [22]. It is harder to separate protein modforms by chromatography but isobaric modforms, such as positional isomers, can be isolated within the spectrometer, thereby simplifying the analysis. Fragmentation (MS*^n^*) can again help to determine modforms but is more difficult to undertake with good coverage for intact proteins. With current TD MS technologies, which rarely go beyond MS^3^, the information shortfall required to determine the modform distribution is reduced but still increases exponentially [3].

An alternative approach to estimating the modform distribution arises from realising that all MS methods lead to linear equations in the modform amounts [3]. For example, suppose that the amounts of modforms 1 to 16 in Fig. 1 are *x*_1_,···*, x*_16_. These are the coordinates of the modform distribution in the 16-dimensional modform space. If BU MS is undertaken with a protease which cleaves between the second and third sites and the modform of the first peptide in which both first and second sites are occupied (blue and magenta colours) is measured, then it follows that,

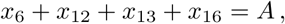

where *A* is the amount of the peptide modform. The protein modforms numbered 6, 12, 13 and 16 contribute the appropriate peptide modform after cleavage, while the other protein modforms do not. Similarly, if we assume that each colour in Fig. 1 represents a PTM with a different mass, then it is possible to determine by TD MS the total amount of those protein positional isomers with, for instance, one blue and one green PTM. This yields the equation,
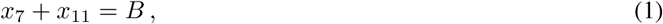

where *B* is the total amount. These linear equations cannot determine the modform region, as noted above, but they do constrain it, especially when taken together with the requirement that amounts cannot be negative, so that *x*_1_,···*,x*_16_ ≥ 0. For instance, this implies from Eq.1 that 0 ≤ *x*_7_ ≤ *B.* Additional equations may constrain this range still further [18]. The totality of equations arising constrain the modform distribution to lie within a bounded region in the high-dimensional space of all modforms (Fig. 1, right). Because of the linearity of the equations, this region must be convex and polyhedral. We refer to it as the “modform region”. The high-dimensional shape of this region can be informative as to which modforms dominate the population. By perturbing the cellular conditions, the change in shape of the region can tell us which PTMs are implicated. It may then become feasible to test how information is being represented by PTMs across the entire protein and to thereby unravel the nature of PTM coding. The modform region can be thought of as a data-centric proxy for the modform distribution.

With that idea in mind, the present paper puts forward a methodology for approximately estimating the shape of the modform region from MS data. It is based on linear programming, which offers an efficient algorithm for determining optimum solutions to linear equations or inequalities. We describe the approach and show how it works with simulated data. This gives the first insights into high-dimensional modform regions. We discuss the problems of using these methods on actual data.

## RESULTS

### Linear equations for MS methods

It is necessary to have a systematic way to generate the linear equations described above. In previous work, we introduced a mathematical formalism for doing so [3] but this was restricted to binary modifications, such as phosphorylation, which are either present or absent. This restriction permits a modform to be identified with the subset of modified sites. Here, we extend the formalism to allow for more complex modifications [19, 27]. We explain the formalism in generality but, for clarity of exposition, focus on those PTMs which are most relevant to the data acquired below. A more complete treatment will be given subsequently. We use set theory notation, as explained in [3], which may be consulted for more background.

Suppose that a protein has *n* sites of modification (hereafter, “sites”), indexed 1, ··· *, n* in order from, say, the N-terminus. Let *S* = {1, ··· *, n*} be the set of sites. Because different PTMs target different amino-acid residues, it is necessary to keep track of which residue occurs at which site. Let 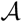 denote the set of relevant amino-acid residues. For instance, if only phosphorylation is being considered, we might take 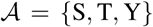, using the customary one-letter codes for amino-acids. We note that phosphorylation may also occur on H and E [19] but ignore that here for simplicity. Let 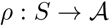 be the residue map, which assigns to each site *i* ∊ *S,* the corresponding amino-acid residue, 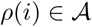. This residue map is a property of the particular protein under study.

We now specify PTMs and define modforms. Let 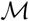 denote the set of relevant, structurally-distinct PTMs, including 0 for the absence of modification. For instance, if acetylation and methylation are being considered, then 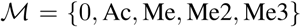, using the customary abbreviations for PTMs. Polymeric modifications, such as ubiquitination and ADP-ribosylation, present a complication, as the number of distinct structures may be unbounded and must be indexed in some manner [19]. In principle, this could be done but the details are beyond the scope of the present paper and we ignore such PTMs here for simplicity. We can think of a protein modform as a function, 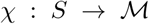, which assigns to a site *i* ∊ *S,* the corresponding PTM, 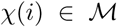. However, such an assignment must be consistent with the residue map *ρ.* The precise consistency requirement will depend on 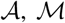, and *ρ* but if we consider as an example phosphorylation, acetylation and methylation, so that 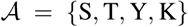 and 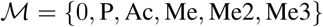, then for 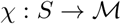 to be consistent as a modform, it is necessary that
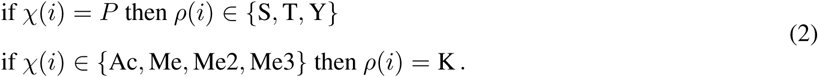

Other consistency conditions can be readily formulated depending on the PTMs being considered. We will say that the function *χ* is consistent and write 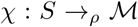 if *χ* satisfies the appropriate consistency conditions with respect to *ρ,* as in Eq.2, for the PTMs under consideration. We can now identify modforms with the consistent *χ*’s. They can be visualised as in the following example modform on 8 sites,
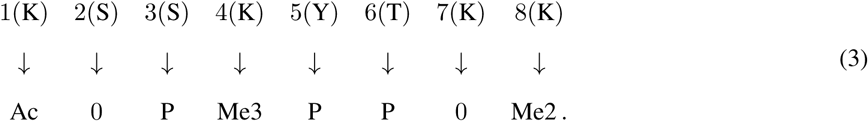

Eq.3 makes clear the resemblance between the modform as defined here and the representation that is often used in the literature, in which the sequence of modifications are listed, K1Ac.S3P.K4Me3.Y5P.T6P.K8Me2. When the sites are known, it is more convenient to denote this Ac.0.P.Me3.P.P.0.Me2 and we will use that format below. The number of potential modforms can be calculated from the consistency conditions. For instance, for the example just considered, the consistency conditions in Eq.2 imply that there are 2 possibilities for the modification state of S, T and Y and 5 possibilities for the modification state of K, so that the total number of combinatorial possibilities is 5 × 2 × 2 × 5 × 2 × 2 × 5 × 5 = 2^4^5^4^ = 10000.

If we only consider a binary modification like phosphorylation, so that 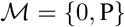, then the function 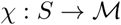 can be identified with the subset of modified sites, {*i* ∊ *S* | *χ*(*i*) = *P*}. Sets were sufficient for our previous work, which involved only such binary modifications [3]. For the more complex modifications considered here, we need functions, 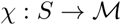.

Let 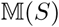 denote the set of protein modforms, 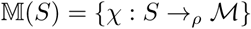. A modform distribution is an assignment to each modform of an amount. This corresponds to a function, 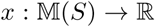, from the set of modforms to the real numbers, 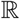. Here, *x*(*χ*) is the amount of modform *χ*, which corresponds to the height of the corresponding bar in the modform histogram in Fig. 1. Since amounts are non-negative, *x*(*χ*) ≥ 0, for all *χ* (which we will write *x* ≥ 0). The functions 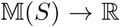 form a vector space, which we will denote 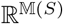 [3]. The dimension of this vector space is given by the number of protein modforms, or the size of 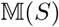. If *X* is any finite set, its size will be denoted *#X*, and we will use *N* to denote the number of protein modforms, so that 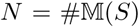. A standard basis for 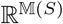 is provided by the unit vectors corresponding to each modform, which lie on the coordinate axes in the modform space in Fig. 1. Let 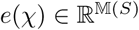 denote the unit vector corresponding to the modform *χ*. As a function on 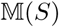,
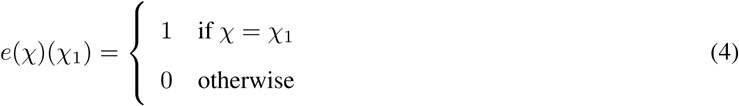

and a modform distribution can be expressed as a linear combination of these basis vectors,

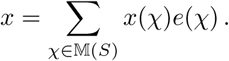

Up to now, we have discussed protein modforms, defined on the entire subset *S* = {1, ···, n}, but the same notation may be used for any segment of the protein that arises through cleavage or fragmentation. Modification sites on the segment are given the same indices as they have in the protein—the protein determines the universe in which the segments are considered—so that a segment can be identified with a subset of sites, *T* ⊆ *S.* This segment has corresponding segment modforms in 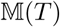. It will be convenient to refer to modforms in 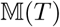 as *T*-modforms, so that protein modforms are *S*-modforms.

Cleavage or fragmentation are two of the basic procedures in mass-spectrometry, out of which many mass-spectrometry experiments are built up. The effect of these procedures is described by a linear segment function, 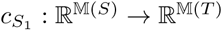, which takes *S*-modforms to *T*-modforms. This function is defined on basis vectors by restriction of the function. If *f* : *X* → *Y* is a function and *X*_1_ ⊆ *X,* then the restriction of *f* to *S*_1_, denoted *f |X*_1_, is just the composition 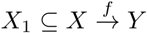. If 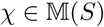, then the segment function is given by,

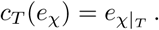

Since this linear function takes a basic vector to a basis vector, its corresponding matrix has entries which are either 0 or 1. Consider, for example, *S* = {1, 2, 3}, *ρ*(1) = *ρ*(2) = S, *ρ*(3) = K, and 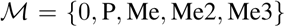. There are 2.2.4 = 16 *S*-modforms. The segment corresponding to the subset *T* = {2, 3} has 8 *T*-modforms. The process of cleavage or fragmentation that creates *T* yields the segment function, *c_T_,* whose 8 × 16 matrix is given by,
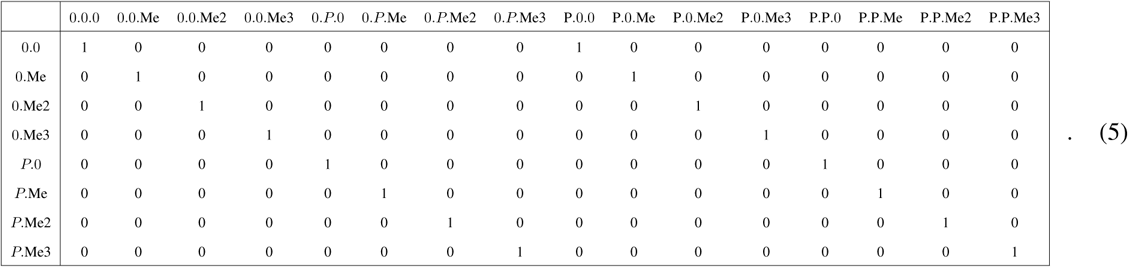

Here, the modforms of the standard basis vectors in 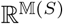 and 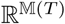 are listed on the top and left, respectively, in the sequence format introduced above.

We have mathematically described cleavage and fragmentation as linear functions on intact proteins, with the domain of the functions being 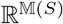. But fragmentation can also be carried out recursively on any segment, *T*_1_ ⊆ *S.* We can define corresponding segment functions on 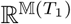 in the following way. Note first that given any pair of subsets, *T*_1_*, T*_2_ ⊆ *S,* for which *T*_2_ ⊆ *T*_1_, there is a natural embedding of the smaller vector space 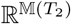 in the larger vector space 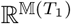. We can consider a *T*_2_-modform, 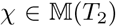, as if it were a *T*_1_-modform by setting all sites in *T*_1_ which are outside *T*_2_ to have no modification. In other words, we define, 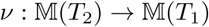 by, for all *i* ∊ *T*_1_,

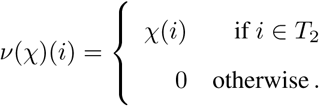

This function 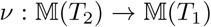 defines an embedding of 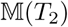 inside 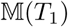. In turn, *v* yields an embedding of 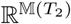 inside 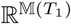, which is defined on basis vectors by sending *e_*χ*_* to *e_v_*_(*χ*)_ for each *T*_2_-modform 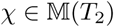. We will denote this embedding, for any pair of subsets, *T*_2_ ⊆ *T*_1_, by 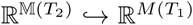. Now, if *T*_2_ ⊆ *T*_1_ is considered to be a fragment of *T*_1_, we can define the *T*_2_-segment function on *T*_1_-modforms by the composition

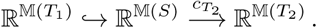

We will denote this composition, with some abuse of notation, also by 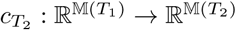.

Cleavage or fragmentation may result in identification and measurement of the segment modforms. This may come about through separation prior to MS or through further fragmentation within the spectrometer or both. In such cases, the end result is an estimate of the modform distribution of the segment, for which matrices like those in Eq.5 give the necessary linear equations. It is also possible that only MS1 is undertaken on the segment. MS1 can resolve modforms with different masses but is unable to resolve isobaric modforms with the same mass. This can be an issue for individual PTMs: the nominal mass of phosphate (80 Da) is the same as that of sulphate and that of acetyl (42 Da) is the same as that of tri-methyl (3 × 14). Modern spectrometers, which are accurate to a few parts per million, can resolve the actual mass differences between such PTMs if the segment is not too large. However, they cannot resolve positional isomers. We will deal with the case of positional isomers here. Extending the formalism to cover isobaric modforms is straightforward but requires more notation.

The target of the resulting linear function is no longer a vector space of the kind 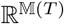. To describe it, let *χ*_1_ ~ *χ*_2_ denote that the modforms 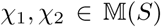 are positional isomers. The precise definition depends on the PTMs involved and care has to be taken with those like methylation which have multiple valencies. If 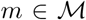 and 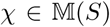, let *n_m_*(*χ*) denote the number of sites having modification *m, n_m_*(*χ*) = #{*i* ∊ *S* | *χ*(*i*) = *m*}. If, for instance, 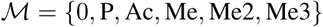, then, *χ*_1_ ~ *χ*_2_ if, and only if,

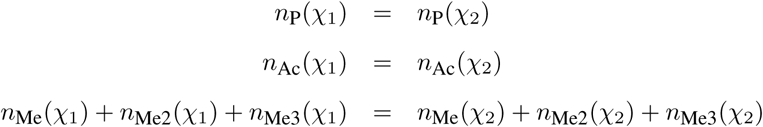

It is clear that the relation ~ is an equivalence relation on modforms and we can therefore form the set of equivalence classes, 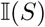. Let 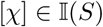 denote the equivalence class containing *χ*, 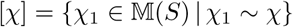, and let *e*_[*χ*]_ be the corresponding standard unit vectors in 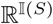, defined in a similar way to Eq.4. Then, MS1 measurement yields a linear mass function, 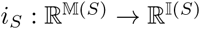, from modforms to positional isomers, which is defined on basis vectors by,

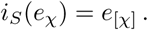

It is straightforward to define positional isomers for any segment *T* ⊆ *S,* which yields the set 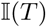 and the corresponding mass function, 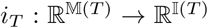. As with cleavage or fragmentation, the resulting matrices, like that in Eq.5, have entries which are 0 or 1.

Mass spectrometry experiments are typically composed of a sequence of the basic procedures of cleavage, fragmentation and MS1 measurement. For instance, the intact protein may be first cleaved by proteolytic digestion into the segment *T*_1_ ⊆ *S,* which is then fragmented into the segment *T*_2_ ⊆ *T*_1_, which is then subjected to MS1 measurement. The overall effect on modforms is described by the linear function which is the composition of the corresponding segment and mass functions,
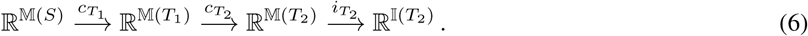

The overall matrix for the composition can be obtained by multiplying the individual matrices. The dimension of the overall matrix in this case is 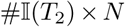, with the number of columns always being the number of protein modforms, as in Eq.5. Since the composition still sends basis vectors to basis vectors, the overall matrix still has entries which are either 0 or 1. The overall matrices arising from each of the individual compositions of basic procedures can be abutted “vertically” to form a single matrix, *M,* of size *r* × *N,* where *r* is the total number of rows of the overall matrices taken together. The matrix *M* summarises the outcome of whatever mass spectrometry experiments have been undertaken as a system of linear equations for the unknown protein modform distribution, 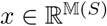,
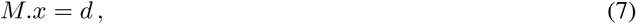

where *d* is the *r* × 1 column vector of actual measurements.

There are other mass spectrometry procedures, such as isolating positional isomers prior to fragmentation [3]. It is not difficult to define linear functions for these, in a similar way to what has been done above, but the procedures described here cover many cases, including those needed for the simulations below. We now turn to asking what can be determined from these linear equations.

### Modform region estimation by linear programming

As shown previously, the system of linear equations given by Eq.7 is not sufficient to determine the unknown modform distribution, 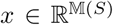, [3] but it can be used to constrain the distribution within a region of 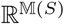 (Fig. 1, right). We can estimate this region by linear programming (LP). LP is about solving (“programming”) the following type of optimisation problem, for which we use the same notation as in Eq.7,
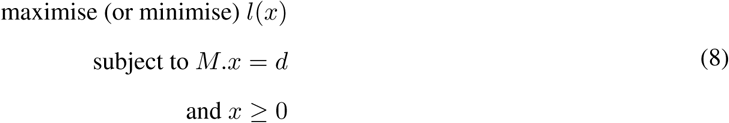

Here, *x* is a *N* × 1 column vector of unknowns, *l*(*x*) is a linear objective function of *x, 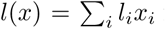* for 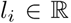, which is specified below, *M* is the known *r* × *N* matrix in Eq.7 and *d* is the *r* × 1 column vector of known data values in Eq.7. Algorithms have been developed which allow LP problems with millions of unknowns to be solved efficiently. This makes LP particularly attractive for modform region estimation, in which the number *N* of unknown modform amounts may be extremely large.

The first requirement for using LP is that the problem should be feasible. In other words, there must be a value of *x* which satisfies the linear system *M.x* = *d.* In our case, the value of *d* will be affected by several kinds of error, arising from sample preparation, instrumentation and measurement, so it is possible that the linear system is infeasible. If so, we first find the smallest perturbation of *d* which yields a feasible solution. The standard procedure is to introduce for each data value, *d_i_,* a pair of non-negative elastic, or slack, variables, *u_i_* ≥ 0 and *vi* ≥ 0, such that the vector of perturbed data values, *d_i_* + *u_i_* − *vi* becomes feasible. We need two non-negative elastic variables because we may sometimes have to increase *d_i_* and sometimes have to decrease it. We can formulate this as an LP problem in which we seek to minimse the total perturbation, 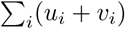, subject to the linear system,

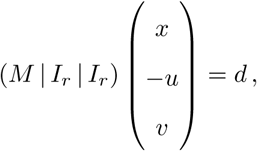

and the inequality constraint *x, u, v* ≥ 0. Here, we have extended *x* “vertically” by abutting the vectors *−u* and *v* to make an unknown vector of size *N* + 2*r* and we have extended *M* “horizontally” by abutting two *r* × *r* identity matrices, to make a matrix of size *r* × (*N* + 2*r*). It is easy to see that this linear system is equivalent to,

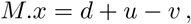

as required. The solution of this LP problem allows us to replace the data vector *d* by the perturbed data vector *d\** = *d+u−v,* for which there is a feasible solution. By minimising the total perturbation, *d\** is the most parsimonious way to reach feasibility, from a linear perspective.

We now want to know the shape of the modform region defined by the feasible linear system *M.x* = *d\**, with *x* ≥ 0. An approximate estimate of the shape can be obtained by using LP to find the minimum and the maximum of each modform amount, *x_i_,*
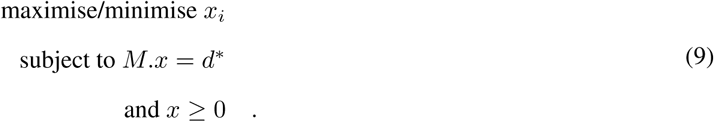

This gives the range within which each modform amount falls, as optimally constrained by the MS data.

Range determination through this LP formulation has the advantage of being easy to undertake efficiently for large *N*. However, it may provide a limited estimate of the actual modform region. For example, if the linear system consists solely of Eq.1 in the Introduction, then the corresponding modform region is the line segment between the points (0, *B*) and (*B,* 0) in Fig. 2. However, the ranges of *x*_7_ and *x*_11_ which come from Eq.9 are both [0, *B*]. This would be same as if the modform region had been the square whose side length is *B* (Fig. 2, magenta box). This is also what happens in general: determination of each range by Eq.9 yields the smallest “hyper-rectangle” whose sides are parallel to the coordinate axes and which contains the modform region (Fig. 2). In the situation in Fig. 2, the hyper-rectangle is a “hyper-square”, with equal sides, but this need not always be the case, as we will see. The individual ranges do not reveal the coupling between *x*_7_ and *x*_11_ which keeps the modform region one-dimensional rather than two-dimensional but its presence can be inferred from the hyper-square structure of the ranges.

**Figure 2:**
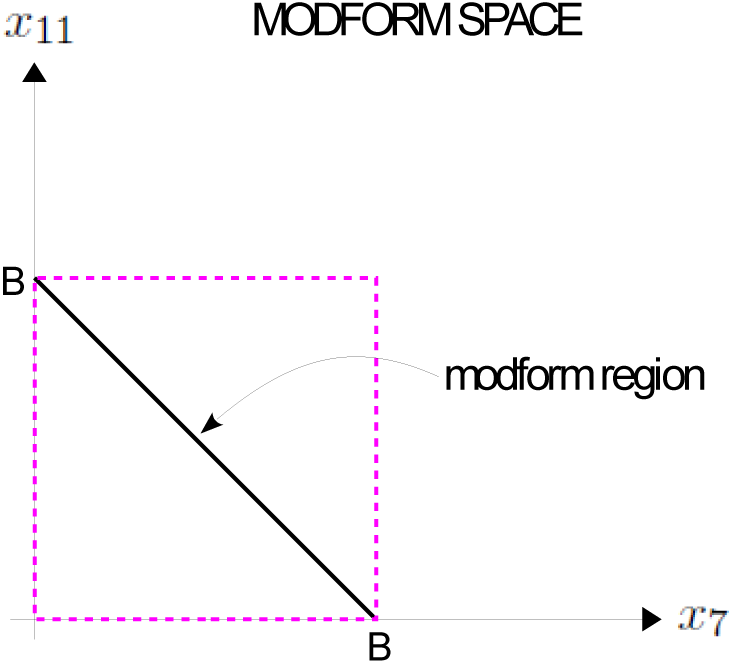
Modform range determination by linear programming (LP). The modform region (black segment) of the linear system given by Eq.7 is shown, in the space of modforms 7 and 11. The ranges obtained by solving the LP problem in Eq.9 gives the same result as if the modform region were the “hyper-square” with magenta dashes.

### Modform regions in high dimensions

We have implemented the LP algorithm in Eq.9 in an open-source software environment, `modformPRO`. This is written in Python and exploits Python’s linear programming library, PuLP (available from https://pythonhosted.org/PuLP/#). The software can take as input MS peak-intensity data, or simulated data, for real proteins, construct the corresponding linear equations for the relevant MS experiments (Eq.7), set up the LP problems (Eq.9), call PuLP to solve them and provide the output as ranges for each modform. We chose as an example the human mitogen-activated protein kinase 1 (MAPK1, Erk1, UniProt ID P28482), with all seven phosphorylations reported on UniProt on the sites S29, T185, Y187, T190, S246, S248 and S284, as marked in Table 1. This gives a total of 2^7^ = 128 modforms. This is well below the software’s capability but our concern in this paper is not with performance of the algorithm but, rather, what it tells us about high-dimensional modform regions, which are investigated here for the first time. In this respect, 128 dimensions is already considerable and the output can only just be visualised on the printed page.

**Table 1:**
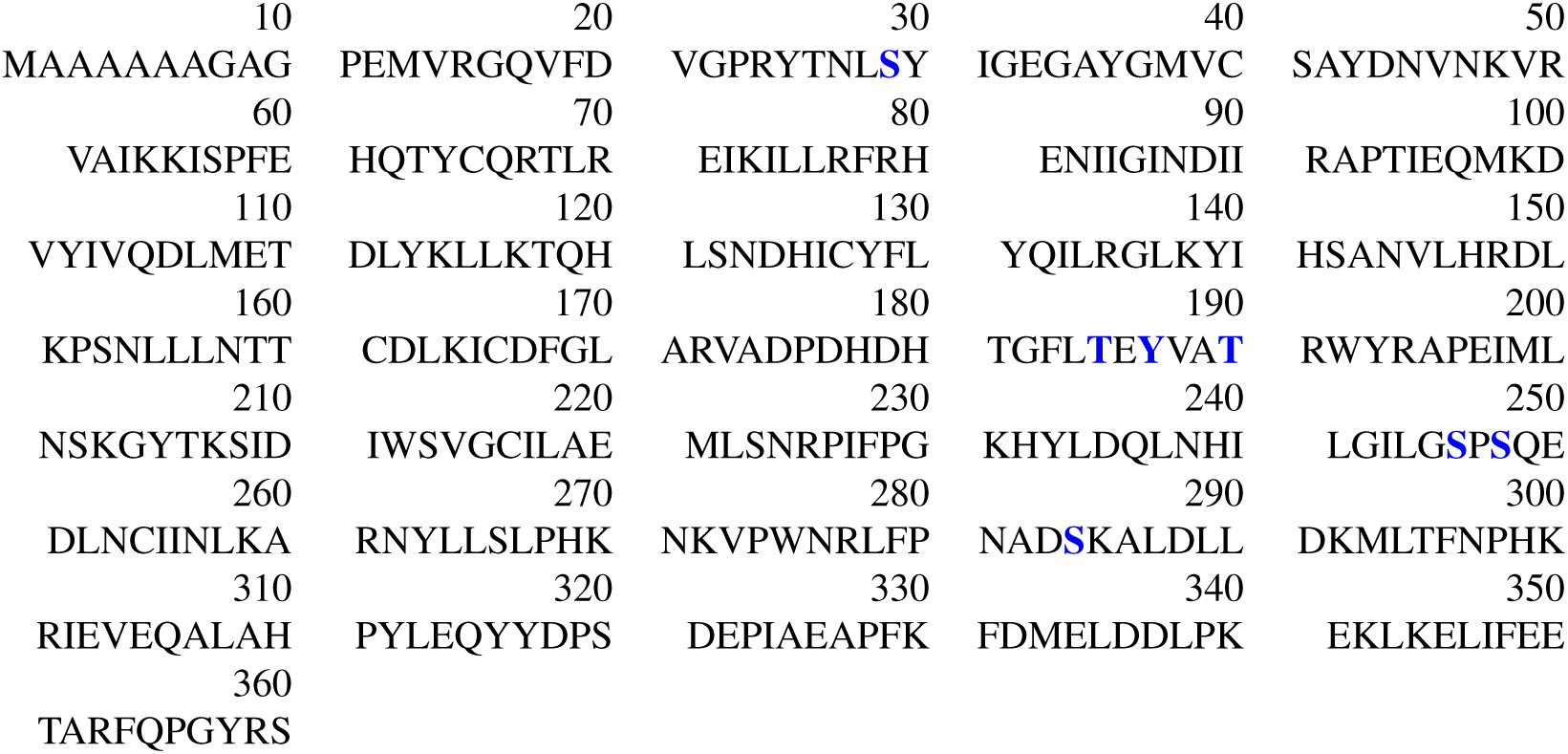
MAPK1 amino-acid sequence for UniProt P28482. The seven phosphorylatable residues annotated in UniProt and used in this study are shown in blue.

We created two simulated modform distributions for MAPK1 as follows. Consider a phospho-modform as a binary string, where 1 marks the presence of P and 0 marks the absence. The Hamming distance between two modforms is then the number of bits by which they differ. The first simulated distribution (“structured”) is one in which the modforms are organised around 4 “modes” with some “noise”. Specifically, we chose 4 modforms at random and gave them each a weight of 100. To each modform at Hamming distance 1 from these 4, of which there are 7, we gave a weight of 10*u* where *u* was a randomly chosen integer between 2 and 8. These are the “modes”. For the “noise”, we chose 20 modforms at random and gave them weights that were randomly chosen real numbers in [0, 30]. If modforms coincided during this procedure, we added up the weights. There should then be nearly 58 modforms with non-zero weights. Finally, we normalised the distribution to the total weight. For the second simulated distribution (“random”), we gave each modform a weight that was a randomly chosen real number in [0,100] and normalised to the total weight.

Although in-vivo data is not yet available, data obtained by in-vitro phosphorylation suggests that modform distributions may be structured, in the sense that few protein modforms arise, despite large numbers of phosphorylated sites [5]. The distinction between the structured and random distributions attempts to reflect this.

In `modformPRO`, we computationally specified one experiment on MAPK1 of BU with tryptic digestion followed by MS1 on the cleavage peptides and one experiment of TD MS1. Fig. 3 shows the range estimations for the structured distribution for each dataset individually and the two datasets combined together. The BU ranges are more variable than the TD ranges, reflecting proteolytic digestion, and they also vary over a broader range, reaching nearly 48% in some cases. However, the average range for BU MS (23.69) is only slightly higher than that for TD MS (22.11). The average range drops much further (16.94) when both datasets are combined.

**Figure 3:**
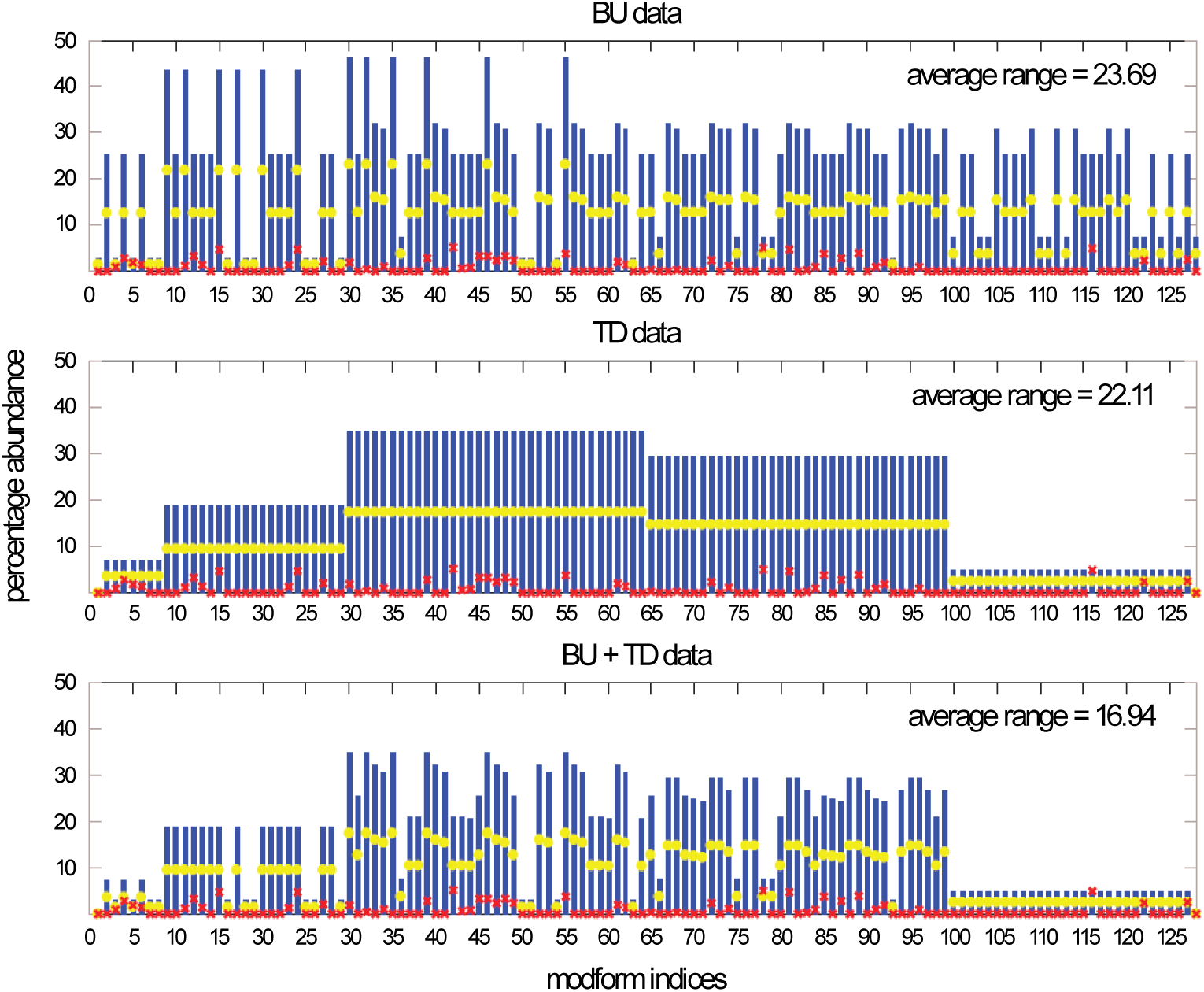
`modformPRO` output for the structured distribution. The modforms corresponding to the indices are shown in Table 2. Red crosses mark the actual values of the modform distribution; blue bars show the range estimated by LP from Eq.9; yellow discs mark the midpoint of each range. The average range is calculated over all 128 modforms.

Fig. 4 shows the range estimation for the random distribution. Since the experiments are the same, a similar pattern of range variation occurs as for the structured distribution. However, in this case the average range for BU (26.76) is considerably higher than for TD (21.04) and the average range for the combined datasets (18.53) does not improve as much over TD.

**Figure 4:**
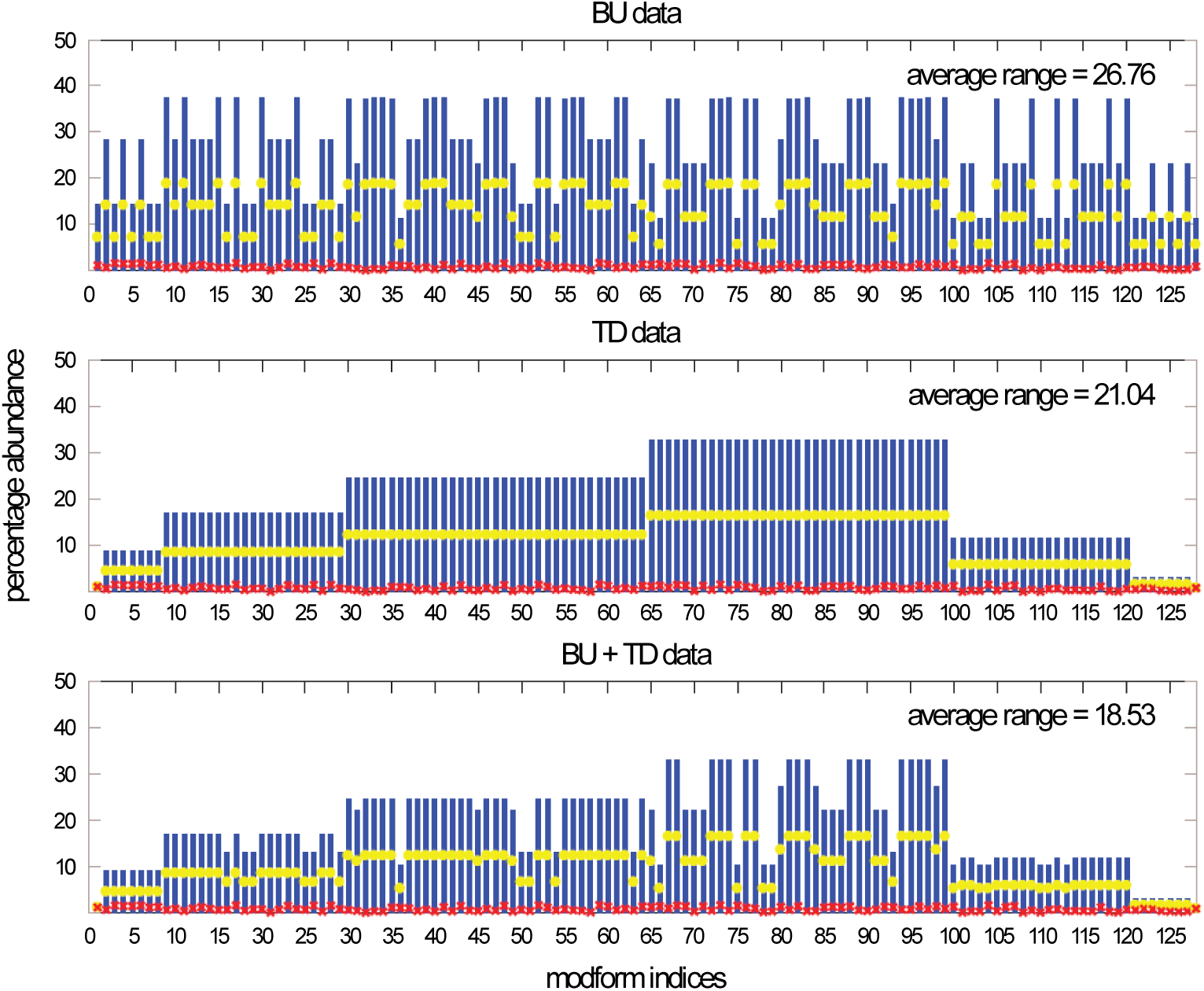
`modformPRO` output for the random distribution, following the same conventions as in Fig. 3. Because of the way weights were chosen for this distribution, as explained in the text, the normalised values (red crosses) must each be below 0.8.

The exact dimension occupied by the modform region depends on the rank of the matrix *M* in Eq.7. The rank is readily determined for TD MS1, as the resulting equations use distinct variables and must necessarily be linearly independent. Here, the rank is 8 and the dimension of the modform region for TD is therefore 120. The rank is visible in the block-like arrangement of the TD plots in Fig. 3 and 4, in which the ranges fall into a small number of distinct sets. TD MS1 cannot distinguish positional isomers, so we expect from Table 2 that the blocks should follow the binomial distribution on 7 sites: 1 (0P), 2 – 8 (1P), 9 – 29 (2P), 30 – 64 (3P), 65 – 99 (4P), 100 – 120 (5P), 121 – 127 (6P) and 128 (7P). Each set of distinct ranges defines a hyper-square like that in Fig. 2. 8 hyper-squares are visible for the random distribution but only 7 for the structured. A closer look at the numerical values shows that the 8th hyper-square is present but is visually indistinguishable in the plot. These hyper-squares reveal the polyhedral shape of the modform region in high-dimensions.

**Table 2:** Indices and modforms for the MAPK1 example. Indices are chosen internally by `modformPRO`.

|  |  |  |  |  |  |
| --- | --- | --- | --- | --- | --- |
| 1 | 0 | 61 | S28P.Y186P.S245P | 121 | S28P.T184P.Y186P.T189P.S245P.S247P |
| 2 | T189P | 62 | Y186P.S245P.S283P | 122 | T184P.Y186P.T189P.S245P.S247P.S283P |
| 3 | S247P | 63 | S28P.S245P.S283P | 123 | S28P.T184P.T189P.S245P.S247P.S283P |
| 4 | T184P | 64 | S28P.Y186P.S283P | 124 | S28P.T184P.Y186P.T189P.S247P.S283P |
| 5 | S245P | 65 | T184P.T189P.S245P.S247P | 125 | S28P.Y186P.T189P.S245P.S247P.S283P |
| 6 | Y186P | 66 | T184P.Y186P.T189P.S247P | 126 | S28P.T184P.Y186P.T189P.S245P.S283P |
| 7 | S28P | 67 | S28P.T184P.T189P.S247P | 127 | S28P.T184P.Y186P.S245P.S247P.S283P |
| 8 | S283P | 68 | T184P.T189P.S247P.S283P | 128 | S28P.T184P.Y186P.T189P.S245P.S247P.S283P |
| 9 | T189P.S247P | 69 | Y186P.T189P.S245P.S247P |  |  |
| 10 | T184P.T189P | 70 | S28P.T189P.S245P.S247P |  |  |
| 11 | T189P.S245P | 71 | T189P.S245P.S247P.S283P |  |  |
| 12 | Y186P.T189P | 72 | S28P.Y186P.T189P.S247P |  |  |
| 13 | S28P.T189P | 73 | Y186P.T189P.S247P.S283P |  |  |
| 14 | T189P.S283P | 74 | S28P.T189P.S247P.S283P |  |  |
| 15 | T184P.S247P | 75 | T184P.Y186P.T189P.S245P |  |  |
| 16 | S245P.S247P | 76 | S28P.T184P.T189P.S245P |  |  |
| 17 | Y186P.S247P | 77 | T184P.T189P.S245P.S283P |  |  |
| 18 | S28P.S247P | 78 | S28P.T184P.Y186P.T189P |  |  |
| 19 | S247P.S283P | 79 | T184P.Y186P.T189P.S283P |  |  |
| 20 | T184P.S245P | 80 | S28P.T184P.T189P.S283P |  |  |
| 21 | T184P.Y186P | 81 | S28P.Y186P.T189P.S245P |  |  |
| 22 | S28P.T184P | 82 | Y186P.T189P.S245P.S283P |  |  |
| 23 | T184P.S283P | 83 | S28P.T189P.S245P.S283P |  |  |
| 24 | Y186P.S245P | 84 | S28P.Y186P.T189P.S283P |  |  |
| 25 | S28P.S245P | 85 | T184P.Y186P.S245P.S247P |  |  |
| 26 | S245P.S283P | 86 | S28P.T184P.S245P.S247P |  |  |
| 27 | S28P.Y186P | 87 | T184P.S245P.S247P.S283P |  |  |
| 28 | Y186P.S283P | 88 | S28P.T184P.Y186P.S247P |  |  |
| 29 | S28P.S283P | 89 | T184P.Y186P.S247P.S283P |  |  |
| 30 | T184P.T189P.S247P | 90 | S28P.T184P.S247P.S283P |  |  |
| 31 | T189P.S245P.S247P | 91 | S28P.Y186P.S245P.S247P |  |  |
| 32 | Y186P.T189P.S247P | 92 | Y186P.S245P.S247P.S283P |  |  |
| 33 | S28P.T189P.S247P | 93 | S28P.S245P.S247P.S283P |  |  |
| 34 | T189P.S247P.S283P | 94 | S28P.Y186P.S247P.S283P |  |  |
| 35 | T184P.T189P.S245P | 95 | S28P.T184P.Y186P.S245P |  |  |
| 36 | T184P.Y186P.T189P | 96 | T184P.Y186P.S245P.S283P |  |  |
| 37 | S28P.T184P.T189P | 97 | S28P.T184P.S245P.S283P |  |  |
| 38 | T184P.T189P.S283P | 98 | S28P.T184P.Y186P.S283P |  |  |
| 39 | Y186P.T189P.S245P | 99 | S28P.Y186P.S245P.S283P |  |  |
| 40 | S28P.T189P.S245P | 100 | T184P.Y186P.T189P.S245P.S247P |  |  |
| 41 | T189P.S245P.S283P | 101 | S28P.T184P.T189P.S245P.S247P |  |  |
| 42 | S28P.Y186P.T189P | 102 | T184P.T189P.S245P.S247P.S283P |  |  |
| 43 | Y186P.T189P.S283P | 103 | S28P.T184P.Y186P.T189P.S247P |  |  |
| 44 | S28P.T189P.S283P | 104 | T184P.Y186P.T189P.S247P.S283P |  |  |
| 45 | T184P.S245P.S247P | 105 | S28P.T184P.T189P.S247P.S283P |  |  |
| 46 | T184P.Y186P.S247P | 106 | S28P.Y186P.T189P.S245P.S247P |  |  |
| 47 | S28P.T184P.S247P | 107 | Y186P.T189P.S245P.S247P.S283P |  |  |
| 48 | T184P.S247P.S283P | 108 | S28P.T189P.S245P.S247P.S283P |  |  |
| 49 | Y186P.S245P.S247P | 109 | S28P.Y186P.T189P.S247P.S283P |  |  |
| 50 | S28P.S245P.S247P | 110 | S28P.T184P.Y186P.T189P.S245P |  |  |
| 51 | S245P.S247P.S283P | 111 | T184P.Y186P.T189P.S245P.S283P |  |  |
| 52 | S28P.Y186P.S247P | 112 | S28P.T184P.T189P.S245P.S283P |  |  |
| 53 | Y186P.S247P.S283P | 113 | S28P.T184P.Y186P.T189P.S283P |  |  |
| 54 | S28P.S247P.S283P | 114 | S28P.Y186P.T189P.S245P.S283P |  |  |
| 55 | T184P.Y186P.S245P | 115 | S28P.T184P.Y186P.S245P.S247P |  |  |
| 56 | S28P.T184P.S245P | 116 | T184P.Y186P.S245P.S247P.S283P |  |  |
| 57 | T184P.S245P.S283P | 117 | S28P.T184P.S245P.S247P.S283P |  |  |
| 58 | S28P.T184P.Y186P | 118 | S28P.T184P.Y186P.S247P.S283P |  |  |
| 59 | T184P.Y186P.S283P | 119 | S28P.Y186P.S245P.S247P.S283P |  |  |
| 60 | S28P.T184P.S283P | 120 | S28P.T184P.Y186P.S245P.S283P |  |  |

The rank for BU MS is more delicate. Each peptide arising from proteolytic cleavage gives rise to linearly independent equations but there are dependencies between the equations from different peptides. We previously determined a formula for the rank for BU MS given any number of proteases and patterns of cleavage [3]. If there is a single protease giving *P* peptides and peptide *i* gives rise to *e_i_* equations, then the rank of the equations coming from all the peptides taken together is,

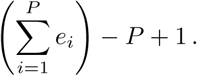

Here, proteolytic cleavage of MAPK1 gives 4 peptides with 1, 2, 1 and 3 modifications and, therefore, 2, 3, 2 and 4 equations, respectively. Hence, the 11 equations arising from BU have rank 11 − 4 + 1 = 8. This modform region for BU therefore also has dimension 120. This is harder to see in the BU range estimation because the hyper-square arrangement is shuffled in the modform ordering. For the structured distribution, 7 hyper-squares are visible in the plot and a closer look at the numerical values reveals an 8th. For the random distribution, 6 hyper-squares are visible and no more are found numerically, presumably because they escape numerical resolution.

No formula currently exists for the rank of the equations for TD and BU combined, although this is work in progress. We independently determined the rank of the 19 equations to be 14, so that the dimension of the modform region decreases to 104. For the random distribution, we found only 12 hyper-squares, suggesting that the remainder were numerically unresolved. However, for the structured distribution we found 17 hyper-squares. These included 2 modforms, the completely unmodified with index 1 and the fully modified with index 128 (Table 2), whose values were exactly determined to be 0. This is not surprising because TD MS1 already accurately accounts for these specific modforms (Fig. 3, middle). The larger number of hyper-squares is unexpected, however. It implies the presence of hyper-rectangles, which are defined by more than one distinct range. This indicates that the polyhedral shape of the modform region has become more intricate. Indeed, the dimensions of the hyper-squares are smaller, and there are more hyper-squares with smaller dimensions, for the structured than for the random distribution (Fig. 5). The smaller the dimension of the hyper-square, the more constrained are the corresponding variables (Fig. 2). We see from Fig. 5 that the polyhedral shape of the structured modform region is more nuanced and considerably more constrained than that of the random distribution.

**Figure 5:**
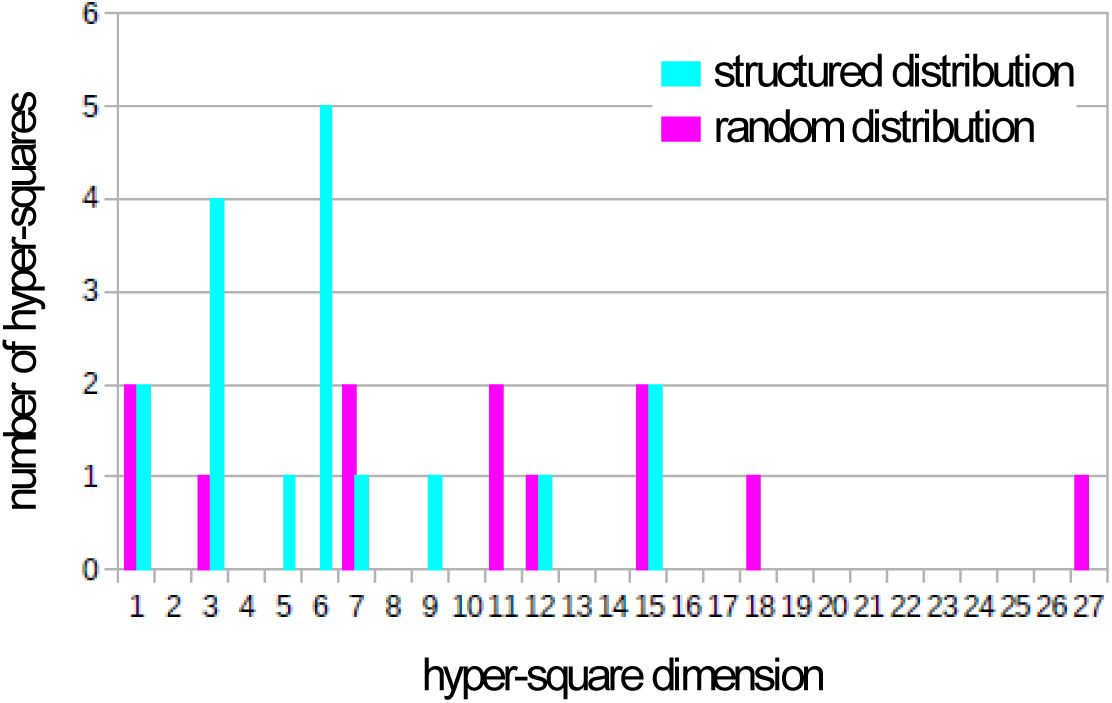
Histogram showing the frequency of hyper-square dimensions arising from combined TD and BU MS, determined from the range estimations for the structured distribution (Fig. 3 bottom plot, cyan) and the random distribution (Fig. 4 bottom plot, magenta).

## DISCUSSION

It is evident from Figs.3 and 4 that the ranges estimated by `modformPRO` are quite coarse and do not constrain the actual modform value tightly. However, the purpose of estimating the modform region is not to recover the modform distribution. This is impossible, as we have discussed. Rather, the modform region is the best that can be done, given the available data. The question is, then, what can be learned about such regions using the LP algorithms implemented in `modformPRO`?

Modforms regions are defined by the linear equations in Eq.7. The more linearly independent equations that are available, the smaller the dimension of the modform region. It is not surprising, therefore, that combining TD and BU data yields a region of smaller dimension. However, the dimension of a region depends only on the matrix *M* in Eq.7 and is the same irrespective of the protein and the modform distribution being analysed. The dimension tells us nothing about the size or shape of the region, which depend both on *M* and on the MS data, which is specified by *d* in Eq.7 and which represents the modform distribution. We have seen that combining TD with BU data reduces the size, as measured by average modform range, for both the structured and the random distributions. This reiterates the importance of combining MS methods, especially combining those which cleave proteins (BU and MD) with those which do not (TD).

The shape of the modform region is also informative. Because modforms regions are defined by linear equations, they are convex and polyhedral but the specific polyhedral shape depends, like the size, on both the matrix *M* and the data *d* in Eq.7. The regions that result from BU or from TD have simple polyhedral shape: up to numerical resolution, they consist of hyper-squares, with the number of these being equal to the dimension. The modform region for combined TD and BU has a more intricate polyhedral shape and is more constrained (Fig. 5), at least for the structured distribution that mimics what might be found in reality [5]. This kind of polyhedral shape is important because it has more degrees of freedom that carry potential information about the modform distribution.

We conclude that the LP methods developed here, while offering a coarse approximation to the modform distribution itself, nevertheless provide useful information about the shape of the modform region.

This brings us to the critical question of how these methods can be deployed in practice, on real data. This raises two kinds of problems. First, it is necessary for different kinds of dataset to be calibrated with each other. Protocols for undertaking such calibration are being developed. Second, and more intractable, is that there is no independent method for determining the modform distribution. Once data become available, we can apply the methods described here to determine a modform region but how can we know that the actual modform distribution lies within it? The polyhedral shape of the regions offers a potential answer to this conundrum. We may not know where the modform distribution lies but we can ask whether perturbations to the modform distribution give rise to correlated changes in the modform region. The experiments necessary to test such correlations will be easier to undertake in vitro, by titrating the levels of modification or demodification enzymes [5]. (Similar perturbations can be carried out in vivo but may have unpredictable effects through indirect or feedback connections within the network of enzymes.) The effect of such perturbations on the modform distribution can be reasonably well predicted and the consequent impact on the modform region calculated. If that change is seen in the data, it would confirm that the shape of the modform region is acting as a proxy for the modform distribution. Such experimental tests are the next step towards practical exploitation of modform regions and the LP algorithms introduced here.

## Author Contributions

DA designed and wrote the software, with the help of RTF, BPE, DL, CJD, PDC and PMT. RTF and BPE implemented the public distribution and undertook the data analysis. GL, NLK and JG conceived the project. JG wrote the paper with the help of all co-authors.

## Acknowledgments

DA, DL and JG were supported NIH R01 GM105375 (to JG, GL and NLK). We are extremely grateful to Felix Wong for assistance with Figs.3 and 4.

